# A novel Mannan-specific chimeric antigen receptor M-CAR redirects T cells to interact with *Candida* spp. hyphae and *Rhizopus oryzae* spores

**DOI:** 10.1101/2024.12.12.628200

**Authors:** Júlia Garcia Guimarães, Gabriela Yamazaki de Campos, Michele Procópio Machado, Patrícia Kellen Martins Oliveira Brito, Thaila Fernanda dos Reis, Gustavo Henrique Goldman, Patricia Vianna Bonini Palma, Thais Fernanda de Campos Fraga-Silva, Daniela Cardoso Umbelino Cavallini, James Venturini, Thiago Aparecido da Silva

**Affiliations:** Department of Cellular and Molecular Biology, Ribeirao Preto Medical School, University of São Paulo, Ribeirão Preto, São Paulo, Brazil; Ribeirão Preto Pharmaceutical Sciences School, University of São Paulo, Ribeirão Preto, São Paulo, Brazil; Center for Cell-Based Therapy, Regional Blood Center of Ribeirão Preto, University of São Paulo, Ribeirão Preto, São Paulo, Brazil; Institute of Biological and Health Sciences, Federal University of Alagoas, Maceió, Alagoas, Brazil; Faculty of Medicine, Federal University of Mato Grosso do Sul, Campo Grande, Mato Grosso do Sul, Brazil; Department of Clinical Analysis, School of Pharmaceutical Sciences in Araraquara, Sao Paulo State University, Araraquara, São Paulo, Brazil

**Keywords:** Chimeric Antigen Receptor, Invasive Fungal Infection, Jurkat cells, Mannan, *Candida* spp., *Rhizopus oryzae*

## Abstract

Invasive fungal infections (IFIs) are responsible for elevated rates of morbidity and mortality, causing around of 1.5 million deaths annually worldwide. One of the main causative agents of IFIs is *Candida albicans*, and non-albicans *Candida* species have emerged as a spreading global public health concernment. Furthermore, COVID-19 has contributed to a boost in the incidence of IFIs, such as mucormycosis, in which *Rhizopus oryzae* is the most prevalent causative agent. The effector host immune response against IFIs depends on the activity of T cells, which are susceptible to the regulatory effects triggered by fungal virulence factors. The fungal cell wall plays a crucial role as a virulence factor, and its remodeling compromises the development of a specific T-cell response. The redirection of Jurkat T cells to target *Candida* spp. by recognizing targets expressed on the fungal cell wall can be facilitated using chimeric antigen receptor (CAR) technology. This study generated an M-CAR that contains an scFv with specificity to α-1,6 mannose backbone of fungal mannan, and the expression of M-CAR on the surface of modified Jurkat cells triggered a strong activation against *Candida albicans* (hyphae form), *Candida tropicalis* (hyphae form), *Candida parapsilosis* (pseudohyphal form), and *Candida glabrata* (yeast form). Moreover, M-CAR Jurkat cells recognized *Rhizopus oryzae* spores, which induced high expression of cell activation markers. Thus, a novel Mannan-specific CAR enabled strong signal transduction in modified Jurkat cells in the presence of *Candida* spp. or *R. oryzae*.

## INTRODUCTION

The Global Action Fund for Fungal Infections (GAFFI) and the World Health Organization (WHO) consider fungi of the genera *Aspergillus*, *Candida*, *Cryptococcus*, *Pneumocystis*, and *Histoplasma* as the main fungi responsible for invasive fungal infections (IFIs) worldwide ^1,2^. The WHO classified fungal species into priority levels based on criteria such as mortality rate, annual incidence, global distribution, and antifungal resistance to improve the development of therapeutic strategies; *Candida albicans, Candida auris, Candida glabrata (Nakaseomyces glabratus*)*, Candida tropicalis*, and *Candida parapsilosis* were placed in the critical and high priority groups ^3^.

Invasive candidiasis (IC) is often referred to as a deep-seated infection in which *Candida* spp. can reach the bloodstream, causing candidemia and affecting various organs and systems. It is estimated to affect more than 250,000 people annually ^4^. The occurrence of candidemia is usually related to extensive colonization at the site of infection, followed by the failure of host immunity, which allows the translocation of *Candida* spp. from their colonization sites to the bloodstream. A major virulence factor common among some *Candida* spp., such as *C. albicans* and *C. tropicalis*, is the morphological plasticity between the unicellular yeast, pseudohyphae and hyphae (multicellular and filamentous) forms. Yeast form disseminates more effectively, and the hyphal form is adapted to invade the tissue and evade the host’s immune response. The yeast-to-hyphal transition presents a differential expression of cell surface components associated with the remodeling of the layers on the cell wall, which favors the subversion of specific host immunity against candidiasis ^5–8^. *Candida* species without a yeast-to-hyphal transition, such as *C. glabrata* and *C. auris*, possess other virulence factors, such as induced endocytosis, that invade and colonize the host ^9–11^.

The vast majority of pathogen-associated molecular patterns (PAMPs) from fungi are present in the cell wall, which comprises an inner layer composed of a meshwork of β-glucans and chitin and an outer layer enriched with mannans to form mannoproteins, lipids, and glycoproteins. Remodeling of the cell wall composition is highly regulated in response to environmental conditions and the host–pathogen relationship ^5,12^, and reorganization of the disposition of layers on the cell wall occurs to mask antigens recognized by PRRs ^5,13,14^. For example, *C. albicans* alters the composition and exposure of surface molecules during the yeast-to-hyphal transition, and the chitin content is slightly higher in the hyphal form ^18–20^. Furthermore, the number of α-1,6 mannose backbone of mannan without side chain substitutions is higher in hyphal-shaped cells, whereas β-1,3-glucan expression is predominant in the yeast form ^5,15^. The masking of the β-glucan layer by mannans compromises the ability of immune cells to recognize and attack the fungal pathogen ^16^.

Recognition of *Candida* spp. by the host immune response depends on the pattern recognition receptors (PRRs) found in phagocytes recognizes PAMPs to initiate an effector immune response against *Candida* spp., and C-type lectin receptors (CLR) are notably in the recognition of *C. albicans* ^17,18^. The importance of CLR-mediated signaling pathway in the immunity against *Candida* spp. is highlighted by the susceptibility to candidiasis in the deficiency of CARD9 (caspase recruitment domain-containing protein 9) that composes the CLR pathway ^19^. Moreover, CARD9 deficiency favored the establishment of mucosal candidiasis and invasive candidiasis, although Dectin-1, Dectin-2, and Dectin-3 acting in the recognition of β-glucans and α-mannans promote the development of Th17 cell for controlling candidiasis ^20^. *Candida*-specific Th17 subset is generated by antigen presentation by dendritic cells, and αβ T cells and γδ T cells producing IL-17A and IL-17F, respectively, contribute to the induction of antimicrobial peptides by epithelial cells to combat *Candida* growth ^21^. In addition, the activation of CD8+ T cells that differentiate into T cytotoxic cells (Tc) is necessary in mounting a protective antifungal response, and Tc cells act mainly through the release of cytotoxic granules that cause damage to the pathogen^18,22^. CD8 T cells contribute in the defense against candidiasis mainly by IL-17 and IFN-γ produced by Tc17 and Tc1 cells, respectively, that induce the phagocytes activation promoting an immunomodulation in the microenvironment required for immune response to combat candidiasis ^22^.

Systemic antifungal therapy is crucial for the survival and reduction in morbidity caused by IFIs, such as invasive candidiasis, and immunotherapy has become an important research field to tackle the resistance mechanisms developed by fungi against antifungal drugs ^23,24^. Immunotherapy using monoclonal antibodies (mAbs) specific to epitopes displayed on the cell wall has been evaluated for the treatment of invasive candidiasis ^25–27^. The development of mAbs specific to the cell wall components of *Candida* spp. has opened new perspectives for the application of chimeric antigen receptor (CAR) technology that redirects T cells to target *Candida* spp. and other fungi that share the same epitope. CAR combines the effector functions of T lymphocytes and the specific antigen recognition capacity of mAbs ^28^. The application of CAR technology beyond cancer has enabled the evaluation of CAR T-cell therapy to combat IFIs caused by *Cryptococcus* spp. or *Aspergillus* spp. ^29–33^.

In this study, we developed and characterized a mannan-specific CAR (M-CAR) that redirects Jurkat cells to target *Candida* spp. M-CAR was generated including a single-chain variable fragment (scFv) derived from a mAb (2DA6 clone) specific to α-1,6 mannose backbone of fungal mannan produced by Burnham et al. ^27^. The functional and structural characterization of the M-CAR construct was carried out in a human T cell line, Jurkat cells, to optimize the evaluation of CAR expression and its capacity to mediate human T cell activation due to the ease of handling transduction and cell expansion steps ^34,35^. Soluble forms of mannan (obtained from *Saccharomyces cerevisiae*) and *Candida* spp. were recognized by M-CAR, resulting in T-cell activation, as validated by the expression of CD69 and the levels of IL-2 produced after co-cultivation with *C. albicans* and *C. tropicalis* hyphae, *C. parapsilosis* pseudohyphae, *C. glabrata* yeast, and *Rhizopus oryzae* spores. Thus, modification of T cells with M-CAR enables their interaction with *Candida* species and *R. oryzae* responsible for systemic fungal infections.

## MATERIALS AND METHODS

### Cell lines and growth conditions

HEK-293T and Jurkat cell lines (clone E6-1) were acquired from the Rio de Janeiro Cell Bank (BCRJ) and were cultured within a humidified 5% CO2 atmosphere at 37° C. HEK-293T cells were maintained in high-glucose DMEM (Dulbecco’s modified Eagle’s medium, Corning) supplemented with 100 mM sodium pyruvate (Gibco), 1% ampicillin/streptomycin (cat. P4333, Sigma-Aldrich), and 10% fetal bovine serum (FBS, Gibco). Jurkat cells were cultured in RPMI (Roswell Park Memorial Institute medium, Thermo Fisher Scientific) supplemented with 100 mM sodium pyruvate, 1% ampicillin/streptomycin and 10% FBS.

### Culture of fungal species

*C. albicans* (ATCC 64548), *C. tropicalis* (H-2747), *C. parapsilosis* (ATCC 90018), *C. auris* (clinical isolates 467, 468, and 469), and *C. glabrata* (H-3479) were cultured overnight in Sabouraud dextrose (SD) medium (cat. K25-1205, Kasvi) with vigorous agitation (150 RPM) at 30 °C for 15 h. Fungal cell suspensions were washed twice with phosphate-buffered saline (PBS) and resuspended in PBS to determine cell concentration using Neubauer chamber, and the yeast-to-hyphae transition was performed with an incubation of 2 × 10^5^ yeast/mL in RPMI supplemented with 10% FBS for 3–4 h. The yeast and hyphal forms were inactivated using heat (70 °C for 1 h).

*R. oryzae* (IAL 3796) was stocked at -80 °C diluted in SD medium with 50% glycerol, and *R. oryzae* was thawed in SD agar (SDA) and incubated at 25 °C for 7–10 d. *R. oryzae* spores were obtained and washed with PBS, and the spore suspension was homogenized using vortex and incubated at RT for 5 min prior to collecting the cell supernatant that was used for washing with PBS. Spores were heat-inactivated (70 °C for 1 h), and cell concentration was determined using a Neubauer chamber.

### Construction of mannan-specific CAR, M-CAR

The antigen-binding domain of a mannan-specific CAR (M-CAR) was derived from an anti-mannan mAb (2DA6 clone) with specificity for α-1,6 mannan attached to the mannan core. The anti-mannan antibody clone 2DA6 was generated by Burnham-Marusich et al. (2018) and its nucleotide sequence encoding the variable domain was used to express an scFv that comprised the antigen recognition domain of M-CAR. The hinge/transmembrane domain of this CAR was the human CD8α molecule (UniProt P01732, position 136–206 aa) and human costimulatory molecule 4-1BB (UniProt Q07011, position 214–255 aa) was the cytoplasmic portion accoupled with a human CD3ζ molecule (UniProt P20963, position 52–164 aa) in the cytoplasmic region. The M-CAR construct had a FLAG-tag (DYKDDDDK) in the N-terminal region in frame with scFv. The nucleotide sequence encoding M-CAR was subcloned into a second-generation lentiviral vector backbone (pLentiCas9-EGFP, GenScript), replacing a Cas9 sequence, and the vector contained the GFP sequence as a reporter. The lentiviral vector backbone containing only the GFP coding sequence, Mock, was adopted for the modification of T cells, and was considered a negative control in the assays performed.

### Prediction of the molecular structure of M-CAR and stability of scFv

The protein structure homology modeling software SWISS-MODEL (https://swissmodel.expasy.org) was used to generate a 3D conformation of the M-CAR domains ^36^. The molecular structure of M-CAR was performed based on the combination of models predicted related to the M-CAR-composing domains, and the analysis of the electrostatic surface of M-CAR was calculated using APBS and displayed in UCSF ChimeraX ^37^. The stability of the scFv of M-CAR was determined using an scFv containing approximately 60% match of the scFv sequence sourced from anti-mannan mAb (2DA6 clone), and the stability was evaluated using SWISS-MODEL. scFv originating from anti-CD19 FMC63 CAR was used for comparison. The stability of the scFv was predicted using Protein-sol software (https://protein-sol.manchester.ac.uk/) in a physiological-chemical environment (ionic strength, 0.15; pH, 7.5) ^38^. The quantification of scFv stability was based on the energy given in Joules per amino acid followed by a normalization with the molecular size of the protein.

### Production of lentiviral particles containing the M-CAR coding sequence

HEK-293T cell line were used to produce lentiviral particles containing the M-CAR construct-encoding plasmid. The cells were cultivated at a concentration of 2.5 × 10^6^ cells/5 mL of fresh medium in 25-cm^2^ cell culture flasks. After 48 h of incubation at 37 °C, the cell supernatant was reduced to 3 mL. Thereafter, co-transfection was performed using the plasmid pMD2.G (VSV-G envelope-expressing plasmid, cat. no. 12259; Addgene) and psPAX2 (second-generation lentiviral packaging plasmid, cat. no. 12260; Addgene) along with the M-CAR coding sequence. Transfection was performed using Lipofectamine 3000 reagent (cat. L3000001, Thermo Fisher Scientific), following the manufacturer’s instructions. Within 24 h of transfection, 2 mL of fresh medium was added to the culture, after 48 and 72 h of transfection the supernatant containing lentiviral particles was collected. Lentiviral particles were centrifuged (300 × g, 10 min, 4 °C) to remove cells and debris, and to concentrate the lentiviral stock the supernatant was incubated with Lenti-X (cat. 631232; Takara Bio USA, Inc.), according to the manufacturer’s instructions. PBS was used to resuspend the lentiviral stock and aliquots were stored at -80 °C until use.

The measurement of lentiviral particle titer was performed by transduction of Jurkat cells through a spinoculation protocol (850 × g, 65 min, RT). For this, Jurkat cells were seeded in 48-well microplates (1 × 10^5^ cells/250 µL) and 72 h after transduction, the percentage of Jurkat cells expressing M-CAR was detected by GFP expression using flow cytometry. The titer was calculated using the formula “titer = {[(%GFP/100) × dilution × plated cells]/final volume}”, and were considered the dilution of lentiviral particles that resulted in between 10% and 30% GFP-positive cells for the titer determination.

### Modification of Jurkat cells to express M-CAR

Jurkat cell transduction was performed using spinoculation protocol (850 × g, 65 min, RT), and 1 × 10^5^ cells/250 µL were distributed in a 48-well microplate. Jurkat cells were modified using multiplicities of infection (MOIs) of 1, 3, 5, and/or 10 lentiviral particles containing M-CAR or Mock coding sequence. After 72 h of transduction, GFP expression was measured using flow cytometry (Guava EasyCyte, Guava Technologies, Millipore) to determine the transduction efficiency. Cells were immediately used in the experiments or incubated at 37 °C for expansion, once 3–5 × 10^5^ cells/mL were seeded.

### Detection of M-CAR on the cell surface

The FLAG-tag (DYKDDDDK) expressed in the N-terminal region of M-CAR was used to detect M-CAR expression on the surface of Jurkat cells. For this, 1 × 10^6^ cells were blocked with 0.1 M PBS supplemented with 0.5% bovine serum albumin (BSA, cat. A7906-500G, Sigma-Aldrich), and after 30 min of incubation at 4 °C, PBS was used to wash the cells, which were incubated with anti-FLAG monoclonal antibody (cat. MA1-91878-HRP, Thermo Fisher Scientific). After 30 min of incubation at 4 °C, cells were washed with PBS and incubated with anti-mouse IgG–biotin antibody produced in goat. Subsequently, the cells were washed again and incubated for 30 min at 4 °C with PE-conjugated streptavidin. Mock cells and non-transduced Jurkat cells were subjected to the same conditions described above and were considered negative controls. Detection of M-CAR on the cell surface was assessed using flow cytometry (Guava EasyCyte, Guava Technologies, Millipore), and data analysis was done using FlowJo™ software (version 10, for Windows; Becton, Dickinson and Company).

### Recognition of mannan by M-CAR cells using flow cytometry and fluorescence microscopy analysis

Mannan was obtained from *S. cerevisiae* (cat. M7504, Sigma-Aldrich) and biotinylated using the EZ-Link Hydrazide-Biotin kit (cat. 21339, Thermo Fisher Scientific) according to the manufacturer’s instructions. M-CAR and Mock cells modified with MOI of 5 and 10, respectively, were previously incubated with PBS/0.5% BSA for 30 min at 4 °C, and then incubated with 2 μg/mL biotinylated mannan for 30 min at 4°C. Next, the cells were washed with PBS and incubated with streptavidin-PE (1:100) for 30 min at 4 °C prior to being subjected to flow cytometry. Fluorescence microscopy was performed by incubation with AlexaFluor^TM^ 594 streptavidin (1:100, cat. S11227, Thermo Fisher Scientific), after incubation with biotinylated mannan. To visualize the interaction between M-CAR cells (GFP) and biotinylated mannan (AlexaFluor^TM^ 594), a fluorescence microscope (Zeiss LSM 780 System Axio Observer) was used, and the images were analyzed using the Fiji software (Image J).

### Measurement of IL-2 levels using ELISA

Jurkat cells modified with M-CAR using MOI of 1, 3, 5, or 10 were seeded in a 96-well microplate at a concentration of 2 × 10^5^ cells/200 µL and incubated with different antigens, as follows: mannan obtained from *S. cerevisiae* (1 or 10 μg/mL, cat. M7504, Sigma-Aldrich); β-1,3-glucan from *Alcaligenes faecalis* called Curdlan (10 μg/mL, cat. tlrl-curd, InvivoGen); β-glucan peptide (BGP) from *Trametes versicolor* (10 μg/mL, cat. tlrl-bgp, InvivoGen), inactivated yeast or hyphae forms of *C. albicans, C. tropicalis, C. parapsilosis*, or *C. glabrata*, inactivated spores of *R. oryzae* (ratio of 1:1 cells to fungi); inactivated mixture of bacterial from the human gut microbiota (VSL#3®) including *Bifidobacterium breve*, *Bifidobacterium lactis* (Previously classified as *B. longum*), *Bifidobacterium lactis* (Previously classified as *B. infantis*), *Lacticaseibacillus paracasei* (Previously classified as *Lactobacillus paracasei*), *Lactobacillus helveticus*, *Lactiplantibacillus plantarum* (Previously classified as *Lactobacillus plantarum*), *Lactobacillus acidophilus* and *Streptococcus thermophilus* (ratio of 1:1 of cells-to-bacteria). After 24 and 48 h of incubation, IL-2 levels in the co-culture supernatant were quantified using ELISA (BD OptiEIA Human IL-2 ELISA kit, BD Biosciences) according to the manufacturer’s instructions. Mock-transduced Jurkat cells were used as negative controls.

### Quantification of activation markers by flow cytometry

Cells modified with M-CAR at an MOI of 1, 3, 5, or 10 were cultured with the ligands described in the section “Measurement of IL-2 levels using ELISA”. After 20 h of incubation, the cells were incubated with PBS/1% BSA for 30 min at 4 °C for blocking unspecific binding and washed with PBS prior to the incubation with PE-conjugated anti-human CD69 antibody (cat. 555531, BD Biosciences, BD PharMingen) for 30 min at 4 °C. After 30 min of incubation, the cells were washed with PBS, and the expression level and positive cells for CD69 were determined using flow cytometry (Guava EasyCyte, Guava Technologies, Millipore), and the data obtained were analyzed using FlowJo™ software (version 10, for Windows; Becton, Dickinson and Company). The expression of CD25, an activation marker, was determined using flow cytometry after incubation with a PE-conjugated anti-human CD25 antibody (cat. 555432, BD Biosciences, BD PharMingen).

### Apoptotic and exhaustion markers for cells modified with M-CAR in the presence of ligands

Apoptosis was determined by labeling the cells with Annexin V PE (cat. 51-65875X, BD Biosciences, BD PharMingen) according to the manufacturer’s instructions. M-CAR cells modified with different MOIs and Mock cells were incubated with heat-inactivated yeast and hyphal forms of *C. albicans* or soluble mannan, and after 24 h of incubation, the Annexin V PE (2 μL/well/200 μL) diluted in binding buffer (0.1 M Hepes pH 7.4, 1.4 M NaCl, 25 mM CaCl_2_) was added. Cells were incubated for 30 min in a humidified atmosphere with 5% CO_2_ at 37 °C, and cells were harvested before being subjected to flow cytometry (Guava EasyCyte, Guava Technologies, Millipore). As_2_O_3_ (12 μM) was used as a positive control for apoptosis induction.

The expression of the exhaustion markers PD-1 and TIM-3 was evaluated using anti-PD1 (cat. 560795, BD Biosciences, BD PharMingen) and anti-TIM3 (cat. 563422, BD Biosciences, BD PharMingen) mAbs, and the percentage of positive cells and expression level for exhaustion markers were analyzed using flow cytometry (Guava EasyCyte, Guava Technologies, Millipore). Modified M-CAR cells with an MOI of 5 were incubated with heat-inactivated yeast or hyphal forms of *C. albicans*, and the expression rate of exhaustion markers was measured after 24 h of co-cultivation with the fungi, and unmodified and Mock cells were used as negative controls. Staining for exhaustion markers was performed as previously reported for the analysis of activation markers by flow cytometry.

### Statistical analysis

Prism 9.0 (GraphPad Software) was used to analyze the data. The Shapiro–Wilk test was used to determine the normality of all statistical parameters. The homogeneity of variance was evaluated using the F-test for two groups, and the Bartlett test for three or more groups. ANOVA (Analysis of variance) was applied to experiments with three or more groups with Gaussian distributions of the samples, whereas Student’s t-tests were applied to assays with two groups. The Kruskal–Wallis test was used for three or more groups, for datasets with a non-normal distribution, and the Mann–Whitney test for two groups. Differences between group-means were assessed using Kruskal–Wallis test followed by Dunn’s multiple comparison test or one-way ANOVA followed by Tukey’s multiple comparison test. Differences were considered statistically significant at p < 0.05, and the results are showed as mean ± standard deviation (SD).

## RESULTS

### M-CAR express a mannan-specific antigen-binding domain that redirected Jurkat cells to recognize the target

A mannan-specific CAR, named M-CAR, was constructed using an scFv with specificity to α-1,6 mannose backbone sourced from the mAb 2DA6 ^27^. Mannan composes the fungal cell wall, which could be a target for designing new treatment approaches. The DNA sequence encoding variable light (VL) and heavy (VH) chains of mAb 2DA6 were synthetized in frame with coding sequence for hinge/transmembrane from CD8 molecule, followed by signal transduction domains originating from costimulatory CD137 molecule and CD3ζ (Figure 1A). The M-CAR-coding sequence was cloned into a lentiviral vector combined with accessory plasmids, and co-transfection of HEK 293T cells was performed to produce lentiviral particles that carry the M-CAR construct coding sequence (Figure 1A). The titer of the lentiviral particles was determined after transduction of Jurkat cells, and the frequency of GFP-positive cells was measured using flow cytometry.

**Figure 1.**
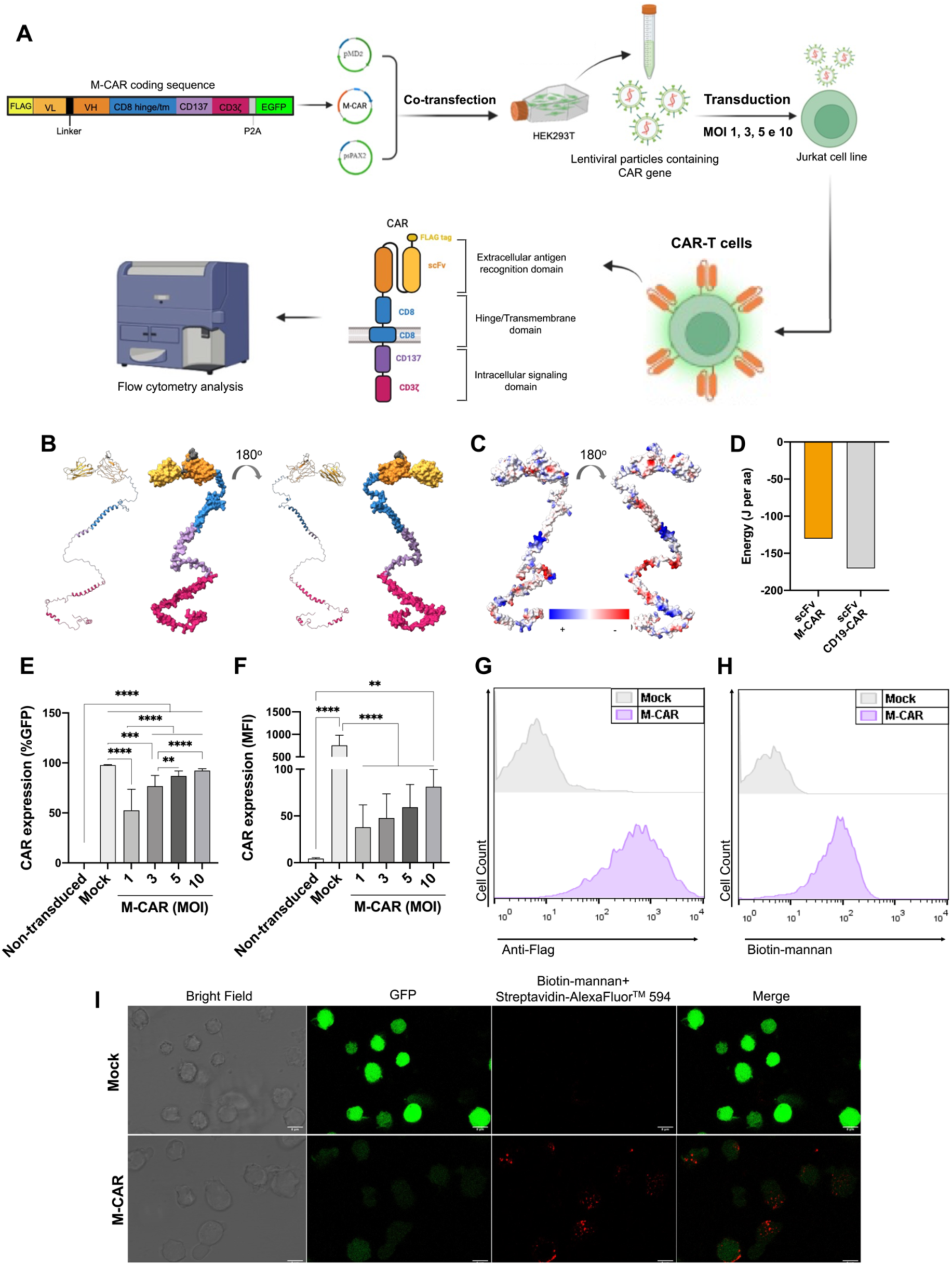
M-CAR-expressing Jurkat cells recognizes mannan. The M-CAR-coding sequence was cloned into a lentiviral vector backbone and used in the co-transfection of HEK-293T cells with accessory plasmids pMD2.G and psPAX2. The cell supernatant containing the lentiviral particles was collected every 24 h for three days, and lentiviral particles were precipitated using a Lenti-X concentrator reagent. The titer of lentiviral particles was determined prior to the modification of Jurkat cells with M-CAR using MOIs of 1, 3, 5, or 10 (A). (B) Computational protein modeling and (C) electrostatic analysis of M-CAR using UCSF ChimeraX. (D) Energies of the scFv of M-CAR and FMC63-scFv of CD19-CAR, at 0.15M ionic strength and pH 7.5, calculated by Protein-Sol web server. Energy is given in Joules per amino acid, followed by normalization with the molecular size of the protein. Three days after transduction, the transduction rate of M-CAR was determined by GFP detection using flow cytometry that was represented by percentage of cells positive for M-CAR (E), and the level of M-CAR expression was represented as mean fluorescence intensity (F). (G) Jurkat cells expressing M-CAR, containing FLAG-tag in the N-terminal, were incubated with murine anti-FLAG monoclonal antibody followed by incubation with goat anti-FLAG IgG biotin-labeled mouse. The detection of FLAG-tag was revealed by streptavidin-PE (x-axis of the histogram). Jurkat cells expressing M-CAR were incubated with 2 μg/mL biotinylated mannan, and the interaction of M-CAR with the target was revealed by streptavidin-PE (x-axis) using flow cytometry (H). (I) Cells modified with M-CAR and Mock were initially incubated with biotinylated mannan and, subsequently, streptavidin conjugated to Alexa Fluor 594 was added. The cells were evaluated by fluorescence microscopy, and M-CAR and Mock cells are visualized in green (GFP) and the recognition of mannan on the cell surface interaction is visualized in red (Alexa Fluor 594). Unmodified Jurkat cells or Mock cells (MOI of 10) were the negative control. Results are expressed as mean ± SD. ** p <0.01, *** p <0.001, **** p <0.0001.

To investigate the molecular structure and biophysical properties of M-CAR, a protein computational modeling technique was applied. Initially, three-dimensional (3D) homology models for each domain of the M-CAR were built using the SWISS Homology Modeler software to create a M-CAR structure (Figure 1B). The electrostatic surface profiles of M-CAR structure performed by UCSF ChimeraX tool that displays negatively charged surfaces in red and positively charged portions in blue (Figure 1C). Furthermore, the molecular stability of the scFv domain from M-CAR was determined using the Protein-Sol server, and the CD19-CAR scFv was adopted as a high stability model (Figure 1D), and the molecular stability between M-CAR and CD19-CAR did not differ significantly.

Initially, Jurkat cells were modified with M-CAR through different MOIs (1, 3, 5, or 10), and the transduction efficiency was determined using flow cytometry (Figure 1A). Jurkat cells modified with M-CAR showed an increase in the expression rate in an MOI-dependent manner, and MOI of 5 and 10 provided high percentages of M-CAR-positive cells (Figure 1E). M-CAR expression was represented by mean fluorescence intensity (MFI) (Figure 1F). Jurkat cells that were not modified with M-CAR and/or those modified with Mock (lentiviral vector backbone carrying only the GFP coding sequence, MOI = 10) were considered negative controls in all assays.

The M-CAR construct expressed a FLAG-tag in the N-terminal region (Figure 1A), which was used to demonstrate M-CAR expression on the cell surface. Jurkat M-CAR cells were incubated with an anti-FLAG mAb, followed by incubation with a PE-conjugated secondary antibody. Jurkat cells modified or not modified with M-CAR were evaluated by flow cytometry, indicating that the majority of M-CAR-expressing cells were positive for the FLAG-tag on the cell surface, whereas Mock cells did not exhibit any anti-FLAG antibody (Figure 1G). As shown in Figure 1H, the ability of M-CAR to recognize the target and M-CAR expression on the cell surface was validated. Mannan obtained from *S. cerevisiae* was biotinylated and incubated with M-CAR Jurkat and Mock cells, and the recognition of biotinylated mannan by M-CAR was revealed by streptavidin-PE conjugation using flow cytometry. This approach demonstrated that M-CAR Jurkat cells recognized soluble mannan, whereas Mock cells did not recognize mannan (Figure 1H).

In addition, fluorescence microscopy was used to visualize mannan recognition by M-CAR. M-CAR Jurkat cells were incubated with biotinylated mannan, followed by incubation with streptavidin-Alexa Fluor 594 conjugate. Visualization of the recognition of mannan by cells expressing M-CAR was performed by fluorescence microscopy, indicating that M-CAR was displayed on the surface of modified cells with an aggregated distribution after mannan recognition (Figure 1I). In contrast, Mock cells incubated with biotinylated-mannan were not revealed with streptavidin-Alexa Fluor™ 594 conjugate (Figure 1I). Taken together, these data indicate that M-CAR is expressed on the cell surface, and that the antigen-binding domain has a high capacity to recognize the ligand.

### M-CAR expression rate decreased during the Jurkat cells expansion, whereas the exhaustion markers did not increase

M-CAR expression on the third-day post-transduction showed transduction efficiency in an MOI-dependent manner, and the effect of M-CAR expression on cellular expansion should be considered over time. M-CAR Jurkat cells modified using MOIs of 1, 3, 5, or 10 were cultivated for 10 d, and the percentage of M-CAR-positive cells and the M-CAR expression rate (mean fluorescence intensity, MFI) were determined by flow cytometry. Then, Jurkat cells previously modified or not (Mock cells) with M-CAR were seeded (4 × 10^4^ cells/well/200 μL) in a 96-well microplate, and cellular expansion and maintenance of M-CAR expression were determined as shown in Figure 2. M-CAR expression using an MOI of 3, 5, and 10 altered cellular expansion compared to Mock cells (Figure 2A), and the concentration of M-CAR positive cells were compromised. The determination of M-CAR+ cells (GFP+) positive for propidium iodide (PI) was performed over time during the cellular expansion, and after 10 days of culture the percentage of non-viable M-CAR+ cells (PI+) was not significantly different compared to Mock cells (Figure 2B). Regarding the maintenance of M-CAR expression over 10 days, the cell viability rate among the groups was established after 10 days of culture, the M-CAR expression rate (MFI) was maintained for each MOI adopted (Figure 2C), the frequency of M-CAR Jurkat cells was maintained at high levels by modification using MOI of 5 and 10 in all periods analyzed (Figure 2D). Then, M-CAR Jurkat cells kept its receptor at high levels of expression without a damage for cell viability, although the cellular expansion rate of M-CAR Jurkat cells was affected by M-CAR expression.

**Figure 2.**
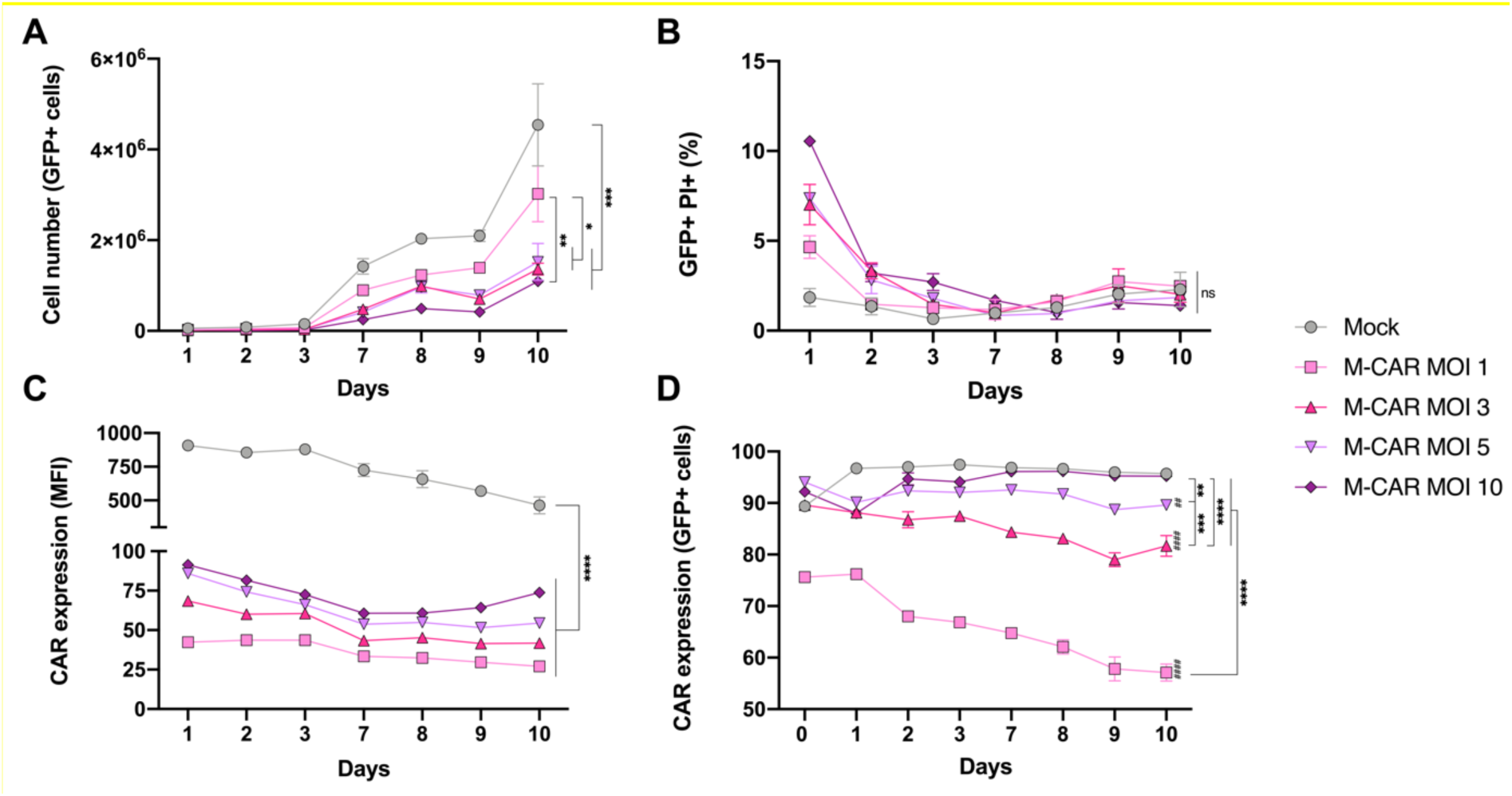
Expansion of M-CAR-modified Jurkat cells maintained the M-CAR expression. Jurkat cells modified with M-CAR (MOIs 1, 3, 5, or 10) or Mock (MOI 10) constructs were distributed in 96-well microplates at a concentration of 2 × 10^5^ cells/mL (200 μL), and cellular expansion and maintenance of M-CAR expression were monitored over 10 d. (A) Cellular expansion represented in absolute number of Jurkat cells modified with Mock or M-CAR (MOI 1, 3, 5, and 10), (B) the cells viability of M-CAR+ cells was evaluated by detection of non-viable cells by propidium iodide (PI) and (C) Mean fluorescence intensity (MFI) of GFP expression, or (D) percentage of cells positive for GFP in Mock or M-CAR cells (MOI 1, 3, 5, and 10). (A-D) The percentage of GFP+ and PI+ cells, the GFP expression rate of the constructs, and concentration of modified Jurkat cells were determined using flow cytometry. # denotes statistical significance between M-CAR and Mock, whereas * denotes statistical significance among the different MOIs used to express M-CAR, as indicated in the graph, and ns denotes non-significant statistical. Results are expressed as mean ± SD. #,* p <0.05; ##,** p <0.01; ###,*** p <0.001; ####,**** p<0.0001.

In addition, Jurkat M-CAR cells modified with MOIs of 3, 5, and 10 did not significantly change the expression of the exhaustion markers PD-1 (programmed cell death 1) and TIM-3 (T cell immunoglobulin 3) compared with unmodified Jurkat cells or Mock cells (Supplementary Figure 1A, B, and D). Also, the expression of CD25, a marker of the high-affinity IL-2 receptor α chain, was evaluated and Jurkat M-CAR cells maintained the absence of CD25 expression such as Mock cells (Supplementary Figure 1C and D).

### M-CAR mediated the recognition of soluble mannan triggering a strong cell activation

M-CAR expression on the surface of modified Jurkat cells allowed us to investigate the responsiveness of cells expressing M-CAR in the presence of the target. The effect of the interaction between M-CAR and the antigen on cell activation was determined by measuring the levels of IL-2 and the expression of the activation marker CD69. Jurkat M-CAR cells modified using different MOIs were incubated with mannan, and after 24 h of incubation, the expression of CD69 increased in M-CAR cells compared to that in Mock cells (Figure 3A). Furthermore, the levels of IL-2 in the cell supernatant were measured, and M-CAR cells produced higher levels of IL-2 in response to different concentrations of mannan after 24 and 48 h of incubation compared to M-CAR Jurkat cells cultured with the medium alone (Figure 3B and C). The levels of IL-2 produced by M-CAR cells incubated with mannan increased in a MOI-dependent manner, demonstrating that a higher M-CAR expression rate induced high levels of cell activation (Figure 3B and C). These findings strongly suggest that recognition of mannan by M-CAR triggers T-cell activation.

**Figure 3.**
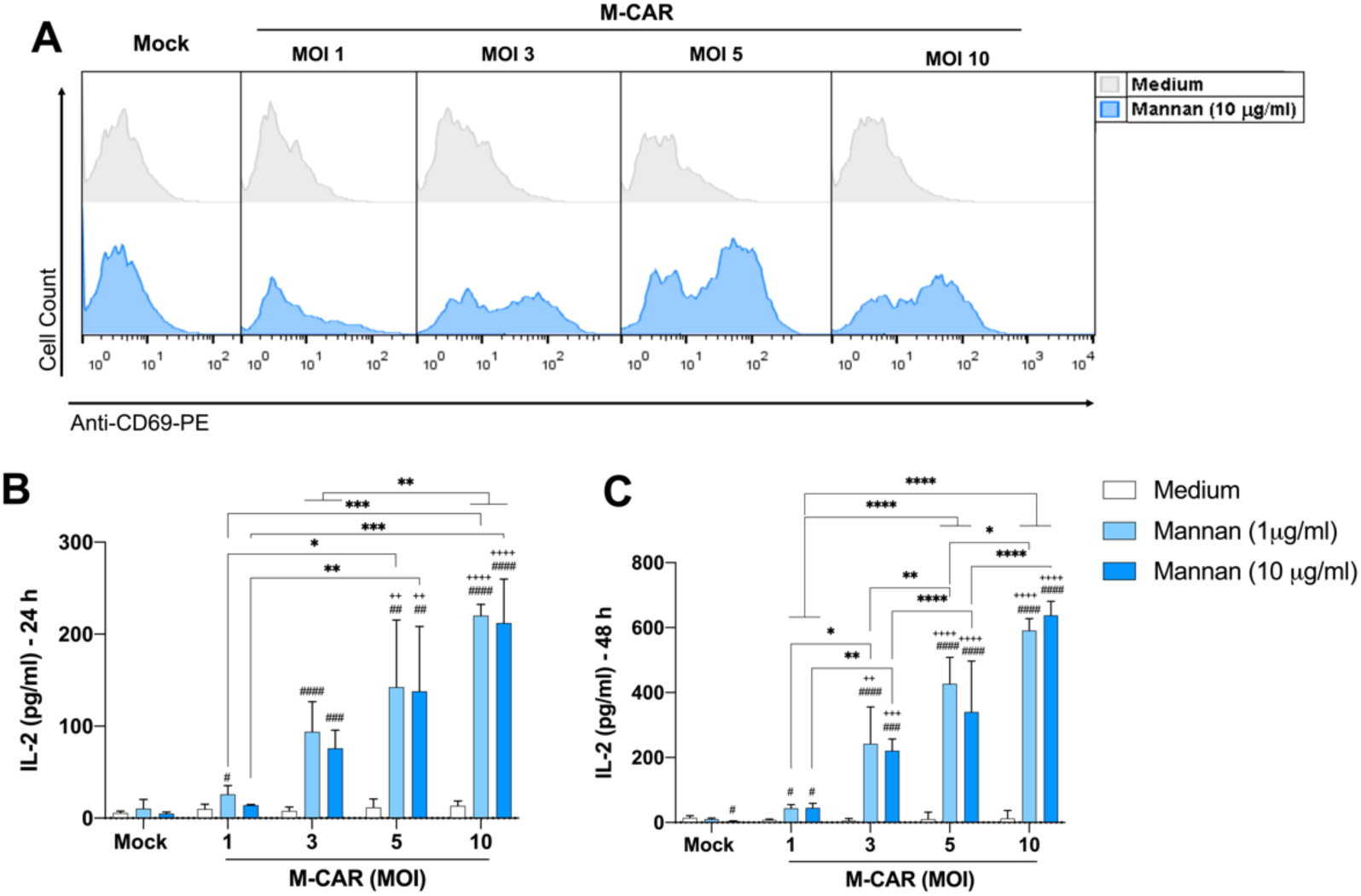
M-CAR induced high expression of CD69 and increased the production of IL-2 in response to incubation with mannan. Jurkat cells were modified with M-CAR using MOIs of 1, 3, 5, or 10, or with Mock using an MOI of 10. Three days post-transduction, the cells were seeded in 96-well microplates (2 × 10^5^ cells/mL) and incubated with mannan (10 μg/mL or 1 μg/mL) from *S. cerevisiae*. (A) After 24 h of incubation, the cells were labeled with monoclonal anti-CD69-PE antibody and detected using flow cytometry. After 24 (B) and 48 h (D) of incubation with mannan, the cell culture supernatant was used to quantify the levels of IL-2 by ELISA. The results are expressed as mean ± SD, and # represents the significances compared to medium within the group and + represents the comparison with Mock. #,*,+ p <0.05; ##,**,++ p <0.01; ###,***,+++ p <0.001; ####,****,++++ p<0.0001.

The specificity of M-CAR was evaluated regarding the ability to induce the cell activation against soluble carbohydrates usually found in the fungal cell wall, such as β-glucan. β-glucan peptide (BGP, extracted from the fungus *Trametes versicolor*) is composed of β-1,4 main chain linked to β-1,3 and β-1,6 glucan side chains, and curdlan (obtained from *Alcaligenes faecalis*) contains β-1,3 glucan. Jurkat cells modified with Mock or M-CAR were incubated for 24 and 48 h with BGP or curdlan, and the CD69 expression rate and the levels of IL-2 released in the cell culture supernatant demonstrated that BGP induced an increase in CD69 expression and IL-2 production in M-CAR Jurkat cells compared to Mock cells (Supplementary Figure 2A-C). The activation markers demonstrated that M-CAR induced slight cell activation against BGP, compared to the cell activation mediated by M-CAR in response to mannan.

### *C. albicans* hyphae was targeted by M-CAR that mediated cell activation

The recognition of soluble mannan by M-CAR, followed by cell activation, allowed us to investigate the ability of M-CAR to redirect modified cells to recognize *C. albicans*. Jurkat M-CAR cells were co-cultured with inactivated yeast and hyphal forms of *C. albicans* (1:1 effector:target ratio) for 24 and 48 h. M-CAR Jurkat cells showed increased expression of CD69 and high levels of IL-2 production after co-cultivation with *C. albicans* hyphae compared with Mock cells or M-CAR cells incubated with the medium alone (Figure 4A-C). The levels of IL-2 produced by the M-CAR Jurkat cells were pronounced in the presence of *C. albicans* hyphae, particularly after 48 h of incubation (Figure 4B and C). Therefore, M-CAR can induce strong T cell activation against *C. albicans* hyphae.

**Figure 4.**
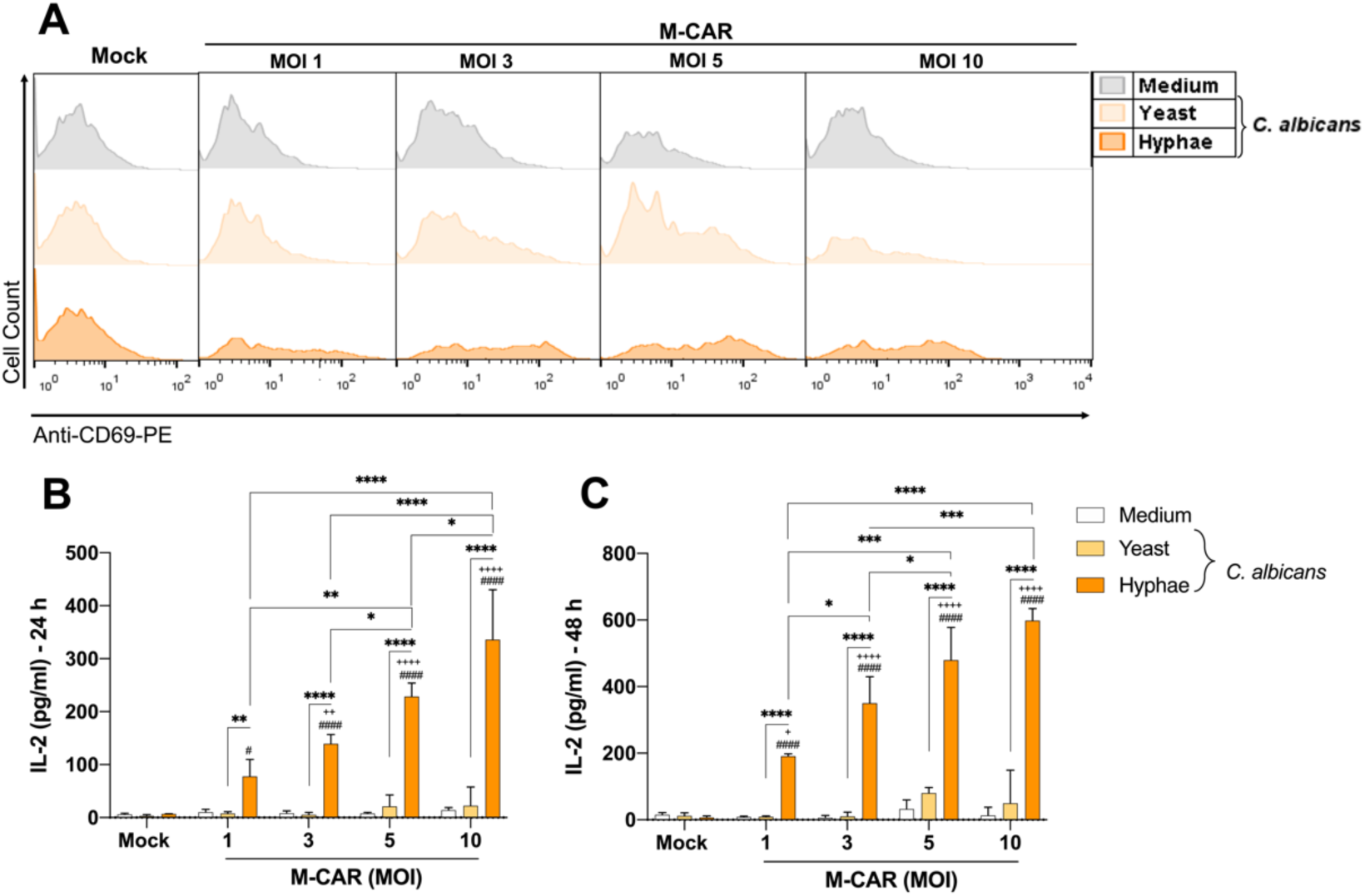
CD69 expression and the IL-2 levels were augmented in the co-culture between M-CAR-expressing cells and hyphae of *C. albicans*. Jurkat cells were modified with M-CAR using MOIs of 1, 3, 5, or 10, and three days post-transduction, those cells were seeded in 96-well microplates (2 × 10^5^ cells/mL). Jurkat cells modified with Mock were the negative control. M-CAR cells were incubated with inactivated forms of yeast or hyphae of *C. albicans* (effector/target ratio = 1:1). (A) After 24 h of co-cultivation, M-CAR cells were incubated with monoclonal anti-CD69-PE antibody and the expression was determined using flow cytometry. (B-C) The levels of IL-2 in the cell culture supernatant were quantified using ELISA. The results are expressed as mean ± SD, and # represents the significances compared to medium within the group and + represents the comparison with Mock. #,*,+ p <0.05; ##,**,++ p <0.01; ###,***,+++ p <0.001; ####,****,++++ p<0.0001.

To evaluate the intensity of M-CAR Jurkat cells against ligands, the expression of exhaustion markers PD-1 and TIM-3 was determined to support full activation mediated by M-CAR against the yeast or hyphal forms of *C. albicans*, as previously demonstrated. Jurkat cells modified with Mock or M-CAR were incubated with yeast or hyphae of *C. albicans* for 24 h, and PD-1 expression increased significantly in M-CAR Jurkat cells co-cultured with *C. albicans* hyphae compared to that in Mock cells or M-CAR cells incubated with medium alone (Figure 5A and C). In contrast, the frequency of M-CAR cells positive for TIM-3 did not alter after incubation with yeast or hyphae of *C. albicans* (Figure 5B and C).

**Figure 5.**
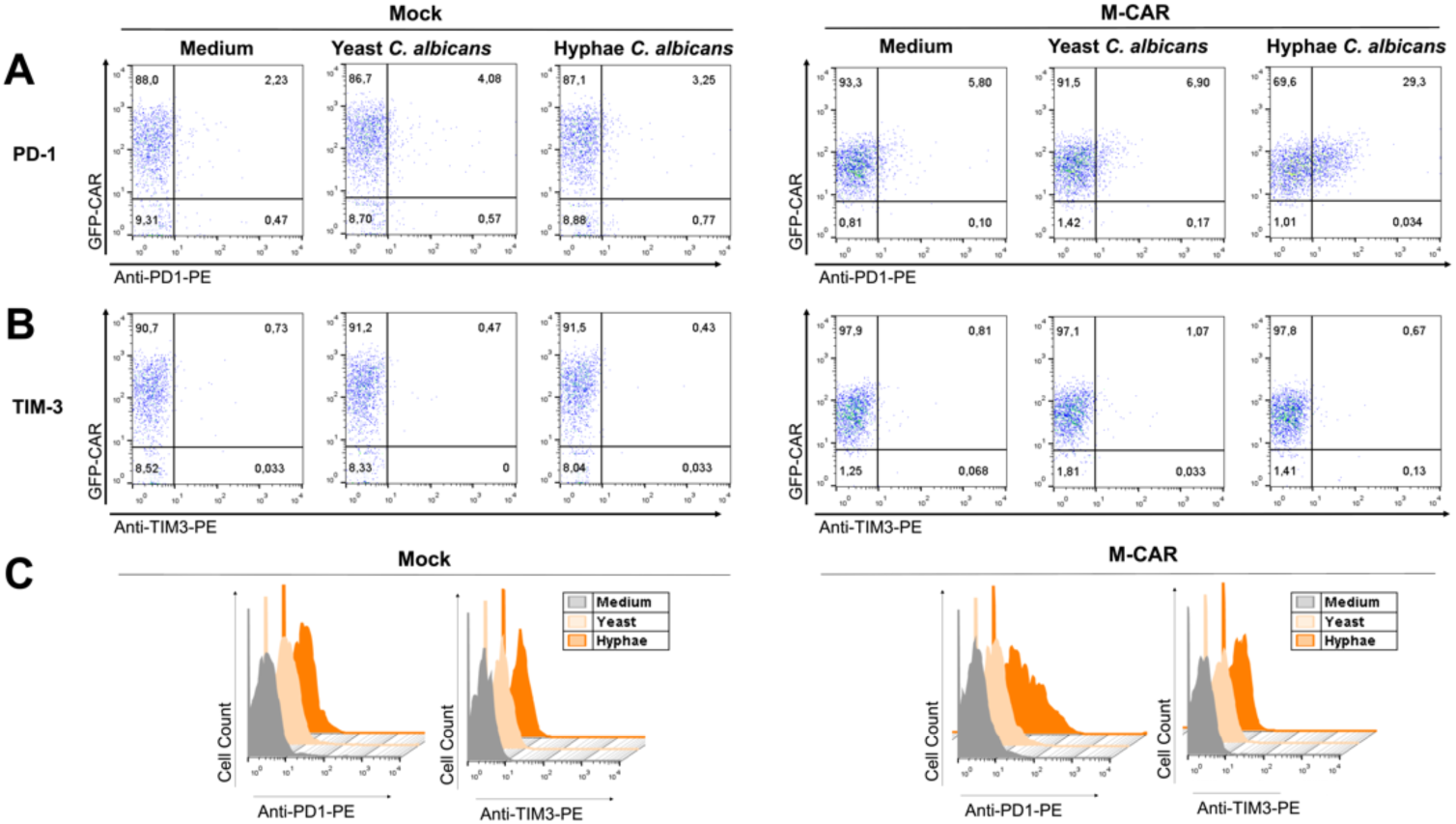
M-CAR-induced cell activation in the presence of hyphal form of *C. albicans* culminated in the expression of exhaustion markers. Jurkat cells modified with Mock (MOI 10) or M-CAR (MOI 5) were incubated with the yeast or hyphal forms of *C. albicans* (effector/target ratio = 1:1), and after 24 h of co-culture, the expression of exhaustion markers was determined using monoclonal antibodies specific to PD-1 (A) or TIM-3 (B). The detection of exhaustion markers was performed using flow cytometry, and the expression of PD-1 and TIM-3 is represented on the x-axis. The expression of M-CAR is observed on the y-axis, and the overlay among the groups is demonstrated by histograms for each marker (D).

The strength of activation of Jurkat cells induced by CAR favors the apoptosis of modified cells, and the detection of annexin V provides an early marker for determining the induction of apoptosis in M-CAR-modified Jurkat cells in response to incubation with mannan or inactivated forms of *C. albicans*. As a positive control for cell death induction, 12 μM As_2_O_3_ was used, which resulted in levels greater than 90% of cells positive for annexin V. M-CAR Jurkat cells were co-cultured with inactivated *C. albicans* yeast and hyphae to analyze the induction of cell apoptosis triggered by cell activation. The frequency (Figure 6A-B) and concentration (Figure 6C-D) of viable and apoptotic cells were determined as follows: (i) percentage of annexin V positive (apoptotic) cells (Figure 6A); (ii) percentage of annexin V positive M-CAR cells (Figure 6B); (iii) concentration of annexin V negative (viable) cells (Figure 6C); (iv) concentration of M-CAR and annexin V negative cells (Figure 6D). Cells modified with M-CAR using MOIs of 1, 3, and 5 and co-cultured with the hyphal form of *C. albicans* or incubated with mannan showed a significant increase in the percentage of annexin V-positive M-CAR cells compared to cells incubated with the medium (Figure 6B). These findings were corroborated by a significant reduction in the concentration of viable M-CAR Jurkat cells (annexin V-negative cells) in the presence of the hyphal form of *C. albicans* or mannan (Figure 6B and D). The induction of apoptosis in M-CAR Jurkat cells in response to ligands supports the capacity of M-CAR to induce signal transduction in the presence of *C. albicans* hyphae.

**Figure 6.**
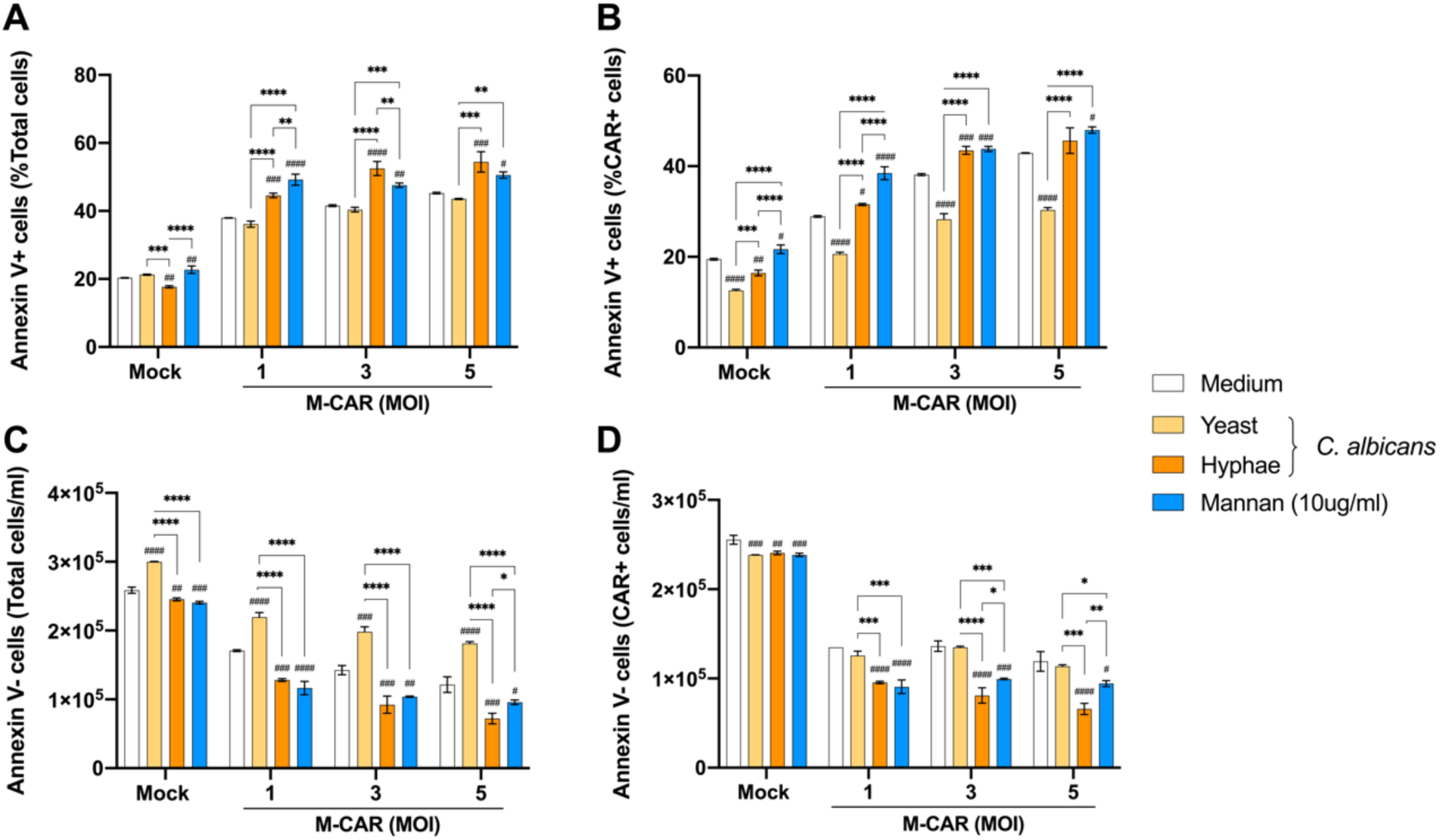
Jurkat cells modified with M-CAR and incubated with *C. albicans* or mannan had high frequency of apoptotic cells. (A-D) Jurkat cells modified with Mock (MOI of 10) or M-CAR (MOI 1, 3, or 5) were incubated with inactivated yeast or hyphal forms of *C. albicans* (ratio of 1:1 of effector to target) or with mannan (10 μg/mL) obtained from *S. cerevisiae*. After 24 h of co-culture, annexin V and binding buffer were added, and after 30 min of incubation the cells were evaluated by flow cytometry. Positive control for induction of apoptosis was considered As_2_O3 (12 μM). (A) Percentage of total annexin V positive cells, (B) percentage of M-CAR annexin V positive cells, (C) concentration of total annexin V negative cells, and (D) concentration of M-CAR annexin V negative cells. The results are expressed as mean ± SD, the significances presented above the bars with “#” are related to the comparison with the mean. #,* p <0.05; ##,** p <0.01; ###,*** p <0.001; ####,**** p<0.0001.

### M-CAR Jurkat cells were activated in response to different *Candida* species and *Rhizopus oryzae*

The light and heavy variable chains sourced from the 2DA6 mAb which was used to compose the antigen-binding domain of M-CAR, were previously demonstrated to recognize mannan from several fungal species such as *Mucor* spp., *Rhizopus* spp., *Aspergillus* spp., *Fusarium* spp., and *Candida* spp.^27^. Once the capacity of M-CAR to induce T-cell activation against *C. albicans* was demonstrated, the interaction of M-CAR Jurkat cells with other fungal species was evaluated. Cells expressing M-CAR were co-cultured with inactivated forms of *C. tropicalis* (yeast and hyphae), *C. parapsilosis* (yeast and pseudohyphae), or *C. glabrata* (yeast). The expression of CD69 on the cell surface and the levels of IL-2 in the cell supernatant were measured after 24 and/or 48 h of incubation. M-CAR Jurkat cells increased the expression of CD69 and showed high levels of IL-2 production in the presence of hyphal and pseudohyphal forms of *C. tropicalis* and *C. parapsilosis*, respectively, and co-culture with *C. glabrata* induced high activation of M-CAR Jurkat cells (Figure 7A-C). Given the broad recognition of M-CAR across clinically relevant *Candida* species, its potential in mediating cell activation against *Candida auris* was also evaluated. *C. auris* is a global concern due to the emergence of multidrug-resistant strains. Cells expressing M-CAR were co-cultured separately with three clinical isolates of *C. auris*, and the measurement of CD69 expression and IL-2 production demonstrated that M-CAR was unable to recognize *C. auris* (Supplementary Figure 3A-C).

**Figure 7.**
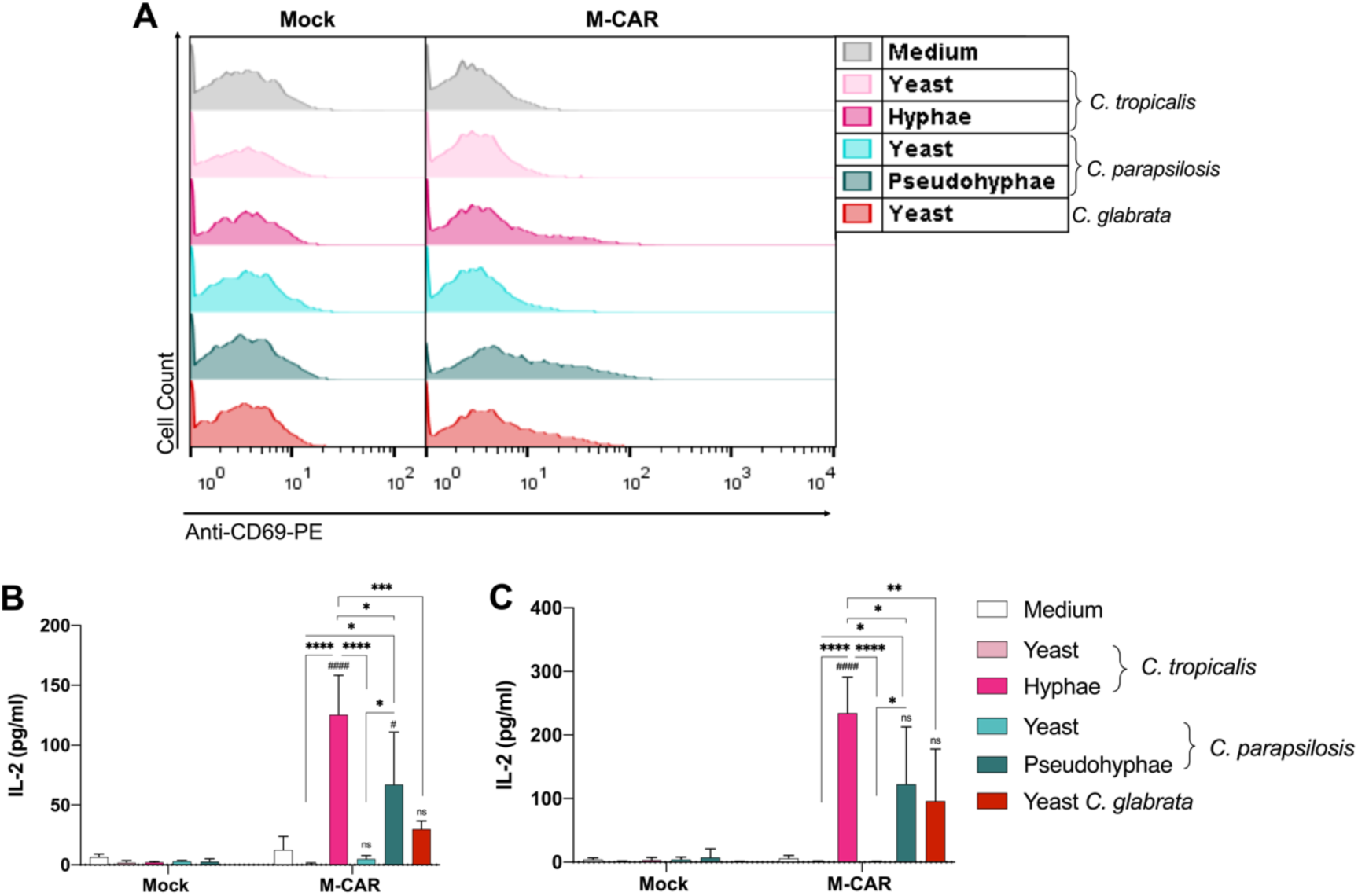
M-CAR induced cell activation against different *Candida* species. (A-C) Jurkat cells modified with M-CAR using MOI 5 and control cells (Mock) were distributed in 96-well microplates at a concentration of 2 × 10^5^ cells/mL and co-cultured with yeast or hyphae forms of *C. tropicalis*, yeast or pseudohyphae of *C. parapsilosis*, and yeast form of *C. glabrata* (effector/target ratio = 1:1). After 24 h of incubation, (A) the levels of CD69 expression were detected using flow cytometry, and after 24 (B) and 48 h (C) of incubation the levels of IL-2 were quantified in the cell supernatant by ELISA. The results are expressed as mean ± SD, and the symbol # represents the significances compared to the medium. #,* p <0.05; ##,** p <0.01; ###,*** p <0.001; ####,**** p<0.0001.

In addition, the ability of M-CAR to induce the activation of modified Jurkat cells in response to *Rhizopus oryzae* was evaluated due to the specificity of mAb 2DA6 for this species^27^. M-CAR cells were co-cultured with inactivated *R. oryzae* spores and CD69 expression was induced within 24 h of incubation with *R. oryzae* (Figure 8A). Moreover, the levels of IL-2 in the cell supernatant increased significantly after 24 and 48 h of co-culture (Figure 8B and C), demonstrating that M-CAR recognizes *R. oryzae* spores and induces cellular activation. The investigation of recognition of other fungal genera and species of clinical importance, such as *C. krusei (Pichia kudriavzevii), C. guillermondii, C. auris, Cryptococcus gatti, Cryptococcus neoformans* and *Aspergillus fumigatus*, was performed through co-culture with M-CAR Jurkat cells, and none of the fungal species considered were recognized by M-CAR, as demonstrated by the levels of IL-2 production (Figure 8D). Thus, the current study characterized the capacity of M-CAR to mediate cell activation against fungi with high clinical relevance. Jurkat cells expressing M-CAR and Mock cells were also co-cultured in the presence of bacterial from human gut microbiota, and the levels of IL-2 production demonstrated that M-CAR did not mediate the T cell activation against the bacterial mixture (Supplementary Figure 4). Furthermore, M-CAR cells showed high levels of IL-2 in the cell supernatant in the presence of *C. albicans* hyphae and bacterial mixture, compared to M-CAR cells in different conditions (Supplementary Figure 4). These findings demonstrated that M-CAR maintained the capacity to target *C. albicans* hyphae even in the presence of gut microbiota indicating that off-target did not occur in this microenvironment.

**Figure 8.**
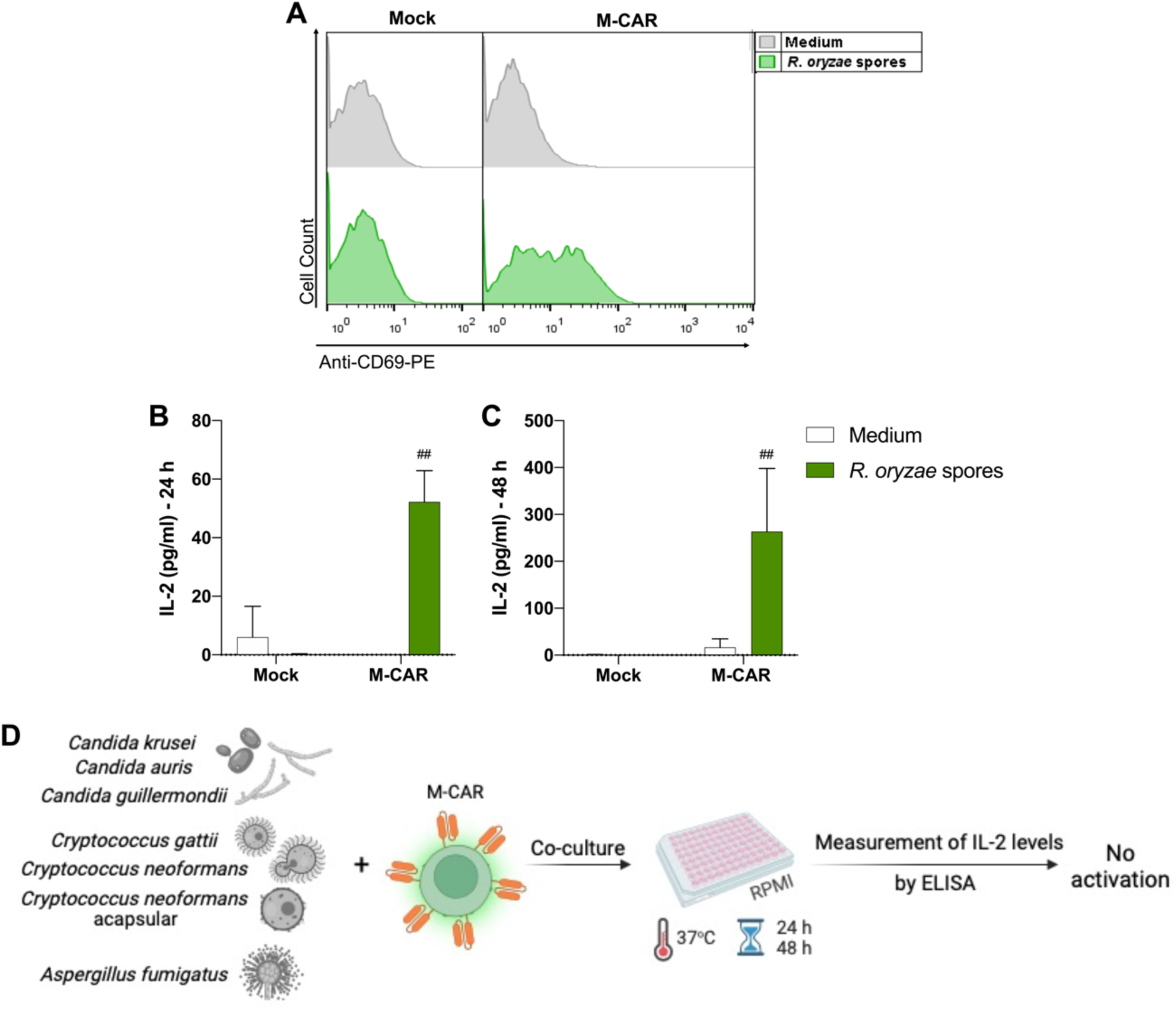
M-CAR-expressing Jurkat cells produced high levels of IL-2 against *R. oryzae*. (A-C) Jurkat cells modified with M-CAR using MOI 5 and control cells (modified with Mock) were seeded in 96-well microplates at a concentration of 2 × 10^5^ cells/mL and co-cultured with *R. oryzae* spores (effector/target ratio = 1:1). After 24 h of incubation, (A) the expression of CD69 was determined using flow cytometry, and after 24 (B) and 48 h (C) of incubation, the levels of IL-2 in the cell culture supernatant were measured using ELISA. (D) M-CAR-expressing Jurkat cells were also incubated with other fungal species and the production of IL-2 was quantified from the cellular supernatant using ELISA after 24 and 48 h of incubation. The results are expressed as mean ± SD, and the symbol # represents the significances compared to the medium. ## p <0.01.

## DISCUSSION

*C. albicans* has been a pathogen with high relevance to the rate of IFIs for several decades, and *Candida* non-*albicans* species have impacted IFIs on a global scale ^39^. The incidence of candidiasis had increased during the COVID-19 pandemic, allowing the occurrence of other opportunistic pathogens such as mucormycosis caused by *R. oryzae* ^40,41^. Invasive fungal infections in CAR T-cell recipients can be triggered by risk factors usually found in hematological malignancies, and candidiasis has been reported ^42^. CAR T-cell therapy against hematological neoplasms can be supported by the application of CAR to redirect T cells to target *Candida* spp. The current work developed a CAR specific to α-1,6 mannose backbone of fungal mannan, named M-CAR, to evaluate the capacity of T cells expressing M-CAR to interact with *Candida* spp. followed by cell activation. M-CAR-modified Jurkat cells were redirected to target *C.albicans and C. tropicalis* hyphae, *C. parapsilosis* pseudohyphae, *C. glabrata* yeast and *R. oryzae* spores, resulting in high IL-2 production and increased CD69 expression. The data evidenced that M-CAR has specificity to α-1,6 mannose backbone exposed by *Candida* species and *R. oryzae* among several fungi species tested and not recognized by M-CAR.

T-cell bioengineering with CAR expression technology has revolutionized cell therapy to treat hematological malignancies, and the redirection of T cells by CAR to target proteins, carbohydrates, lipids, or other molecules has enabled the application of CAR beyond cancer, as initiated by infectious diseases ^43,44^. CAR-modified cells are a promising therapeutic strategy against IFIs, as evidenced in preclinical studies involving *Cryptococcus* spp. and *Aspergillus* spp. infections ^29–33^. Therefore, the development of a new CAR that targets distinct fungal species is essential, and the functional activity of CAR constructs against antigens must be reported. The present study characterized a novel CAR specific to mannan, a component usually found in the fungal cell wall. A mAb sourced from the 2DA6 clone was used to construct M-CAR-containing scFv specific to α,1-6 mannan backbone of fungal mannan. Remarkably, anti-α,1-6 mannan antibody showed strong reactivity against purified mannan and cell extracts from different fungi, such as *C. albicans* and *R. oryzae* ^27^. M-CAR+ cells bind to the target mannan obtained from *S. cerevisiae*, an important carbohydrate presents in the cell wall of several fungi ^5,13,15,45^. In addition, the hinge/transmembrane and signal transduction domains of CAR directly affect its expression, and CD8 molecule as a hinge/transmembrane followed by costimulatory molecule CD137 coupled with CD3ζ facilitated the activation and expression of CAR specific to fungi ^31^. Therefore, the original construct of M-CAR was composed of CD8 molecule as hinge/transmembrane domain and the costimulatory molecule CD137 coupled with CD3ζ to trigger the signaling transduction. Modification of the intracellular domains of M-CAR using distinct costimulatory molecules is in progress to evaluate the improvement in activation induced by M-CAR.

Recognition of soluble mannan by M-CAR induced cell activation, as evidenced by the increased expression of CD69 and high levels of IL-2. mAb 2DA6 recognized the epitope located in the elongation of α-1,6 mannose of mannan, showing a high affinity toward mannans purified from the fungal cell wall enriched with α-1,6 mannose structure, such as *S. cerevisiae, C. albicans*, and *Rhizopus* spp. ^27^. The specificity of mAb 2DA6 was demonstrated in detail using wild yeast or *S. cerevisiae* mutant for Mnn2, which is an enzyme associated with the addition of α-1,2-mannose branching to the α-1,6-mannan backbone, and the absence of Mnn2 did not compromise the high affinity of mAb 2DA6. However, the *S. cerevisiae* yeast mutant for Mnn9, which has α-1,6 mannosyltransferase activity, was unable to form a long mannan backbone with α-1,6 type linkage and, therefore, the interaction with mannan by 2DA6 was extremely committed ^27^. The present study demonstrated that the antigen-binding domain originating from mAb 2DA6, which was considered in the assembly of scFv, maintained its interaction with mannan, promoting M-CAR cell activation against soluble mannan and *Candida* spp. Notably, M-CAR cells were activated in the presence of BGP (peptide-β-glucan) extracted from *T. versicolor* containing a branched polypeptide moiety with a β-1,4 glucan main chain and a β-1,3 glucan side chain followed by branching of the β-1,6 glucan type. This cross-reactivity of M-CAR to BGP can result in a conformational change in the 2DA6 scFv, and further studies must be performed; however, the level of activation triggered by M-CAR against BGP was lower than the levels of activation markers in response to incubation with mannan. M-CAR cells were not responsive after incubation with curdlan, which is a high molecular weight β-1,3-glucan polymer. Therefore, M-CAR induced high cell activation against mannan, whereas incubation with BGP promoted weak M-CAR cell activation. In addition, potential off-target interaction due to the M-CAR recognition to α-1,6-mannan backbone should be evaluated in the human gut microbiota, and the capacity of bacterial mixture from the human gut microbiota ^46^ (containing *Bifidobacterium breve, B. lactis, Lactobacillus paracasei, L. helveticus, L. plantarum, L. acidophilus* and *Streptococcus thermophilus*) to induce the activation of M-CAR Jurkat cells was analyzed. M-CAR cells were not activated in response to the bacterial mixture, and interestingly the ability of M-CAR to mediate the Jurkat cell activation against *C. albicans* hyphae occurred even in the presence of bacterial mixture from the human gut microbiota. Despite of the similarities between the *C. albicans* and bacterial cell walls ^47^, M-CAR kept the specificity to recognize *C. albicans* hyphae, which suggested no off-target in the microenvironment of human gut microbiota.

Remarkably, M-CAR recognized different species of *Candida*, and increased levels of CD69 expression and IL-2 production by M-CAR Jurkat cells indicated that the hyphal forms of *C. albicans* and *C. tropicalis*, pseudohyphae of *C. parapsilosis* and yeasts of *C. glabrata* were the major targets of M-CAR. The levels of CD69 expression and IL-2 production detected in M-CAR cells were different among the *Candida* species co-cultivated, which can be explained by the mannose branch in the mannan backbone, which is distinct in *Candida* species ^48–51^. The α-1,6 mannan form is more exposed in the cell wall of *C. albicans* after the remodeling that occurs in the transition from yeast to hyphae ^5,15^, which is associated with the high levels of IL-2 produced by M-CAR cells against *C. albicans* hyphae. In addition, M-CAR cells incubated with *C. albicans* hyphae showed an increase in PD-1 expression, indicating a cell exhaustion profile, and the association of PD-1 expression with reduced survival of activated T cells is known ^52^. Incubation of M-CAR cells with *C. albicans* hyphae or mannan induced cell activation with an increase in the number of apoptotic cells, as confirmed by annexin V. However, the results obtained with Jurkat cells modified with M-CAR and cultured in the absence of ligands did not show high levels of PD-1 and TIM-3 expression.

The present study also evaluated the recognition of other fungal species by M-CAR cells and found that the distribution of mannan on the cell wall was crucial for the ability of M-CAR to target fungi. The *A. fumigatus* extract was recognized by mAb 2DA6; however, the redirection of M-CAR cells to recognize *A. fumigatus* spores was not performed. *A. fumigatus* has linear mannan chains composed of α-1,6 and α-1,2 mannose linkage and there is an elongation of mannan chain with galactofuran side chains that are covalently bond with inner layers of β-glucan and chitin. The structural organization of the cell wall of *A. fumigatus* differs from that of mannans from *S. cerevisiae* and *C. albicans* ^53^. In addition, *Cryptococcus* spp. was not recognized by M-CAR cells; in this case, the presence of a polysaccharide capsule masked the mannoprotein layer, which is a potential site of interaction for M-CAR^5^.

The tonic signaling mechanism is an important feature that can be observed in CAR-expressing cells; however, this phenomenon was not detected in M-CAR cells, as determined by the lower levels of CD-69 expression and IL-2 production when the M-CAR cells were incubated without the ligand. Several studies have investigated the domains and structural properties of CAR that influence the strength of tonic signaling, which can be affected by CAR oligomerization due to scFv. Cellular functionality, perhaps tonic signaling, can benefit or impair cell activation in response to the target under study ^54–56^. Oligomerization mediated by scFv expressed in the CAR promotes clustering on the cell surface, whereas a homogeneous distribution of CAR is expected in the absence of tonic signaling. Usually, the clustering of CAR is triggered by the intrinsic instability of scFv ^55,57^; however, other research groups have demonstrated that CAR clustering depends on the charge distribution of scFv and that positively charged residues induce spontaneous oligomerization. This hypothesis was corroborated by determining the electrostatic properties of different CARs and identifying more positively charged residues in the composition of scFv belonging to the CAR with strong tonic signaling compared to those receptors with weak tonic signaling ^58^. To investigate the molecular structure and biophysical properties of M-CAR, we employed computational protein modeling. Initially, we built three-dimensional (3D) homology models for each M-CAR domain using SWISS-MODEL, an automated protein structure homology-modeling server. The 3D models were combined to construct a complete M-CAR structure. This modeling was subjected to surface electrostatic profile analysis using the UCSF ChimeraX tool, which displayed negatively charged surfaces in red and positively charged portions in blue. The molecular stability of the M-CAR scFv domain was evaluated using the Protein-Sol server. A similar level of stability was achieved between M-CAR and CD19-CAR scFv, which did not present tonic signaling ^59^. Therefore, our results related to the detection of a balanced distribution of positively and negatively charged residues in M-CAR align with the description of the absence of tonic signaling in cells expressing M-CAR.

## CONCLUSION

Jurkat cells modified with M-CAR showed a high capacity to recognize mannan, α-1,6 mannose backbone, localized in the cell wall of *C. albicans* hyphae and *R. oryzae*. Cellular activation markers demonstrated that M-CAR induced strong signal transduction in modified Jurkat cells in the presence of *Candida* spp. and *R. oryzae*.

## ACKNOWLEDGEMENT

We thank Prof. Maria Cristina Roque-Barreira and Prof. Paulo Coelho (Ribeirão Preto Medical School, University of São Paulo, Ribeirão Preto, Brazil) for providing infra-structure support.

## AUTHORS CONTRIBUTION

Conceptualization, J.G.G and T.A.S; methodology, J.G.G, G.Y.C, M.P.M, P.K.M.O.B, T. F. R, G. H. G, T. F. C. F. S., J. V., P. V. B. P., D. C. U. C. and T.A.S; software, J.G.G, G.Y.C, M.P.M, P.K.M.O.B, and T.A.S; writing – original draft preparation, J.G.G and T.A.S; writing – review and editing, J.G.G, G.Y.C and T.A.S; supervision, T.A.S.; funding acquisition, T.A.S. All authors have read and agreed to the published version of the manuscript.

## DATA AVAILABILITY STATEMENT

The data that support the findings of this study are available from the corresponding author, da Silva TA, upon reasonable request.

