## Supplementary Material for "A novel Mannan-specific chimeric antigen receptor M-CAR redirects T cells to interact with *Candida* spp. hyphae and *Rhizopus oryzae* spores"

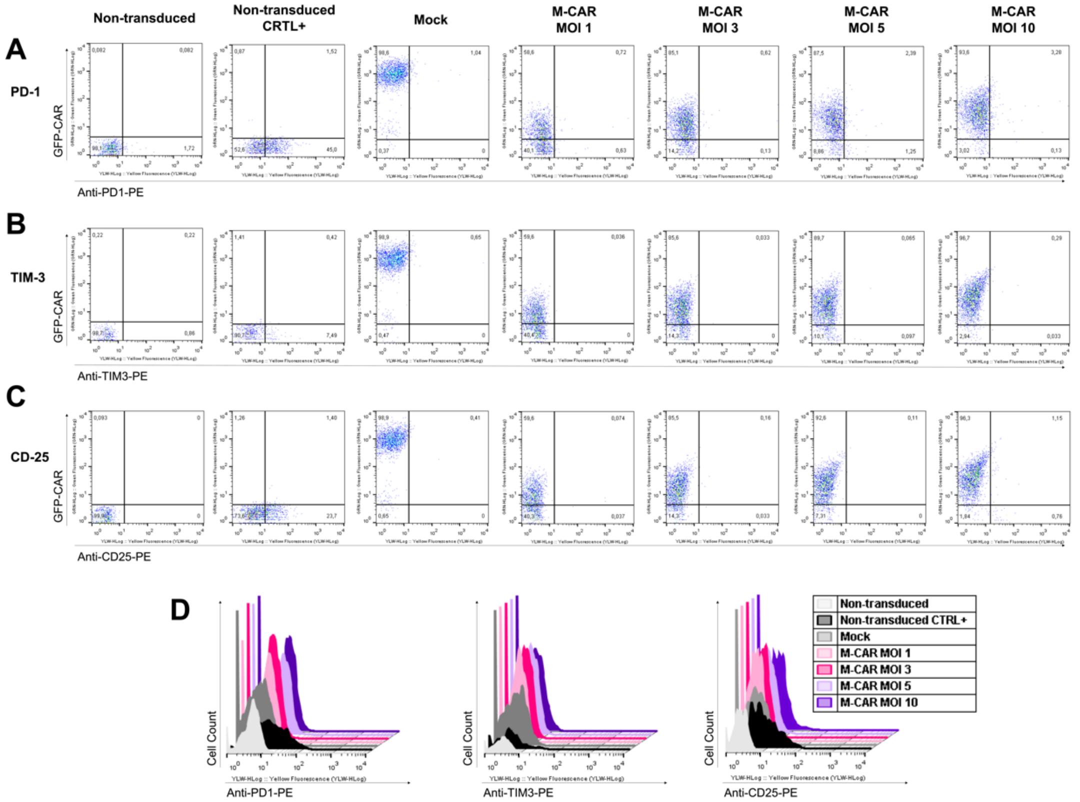


**Supplementary figure 1. Surface markers expressed in Jurkat cells modified with M-CAR from different MOIs.** Jurkat cells were transduced with M-CAR at MOIs of 1, 3, 5, and 10, and unmodified cells or cells transduced with Lenti-mock at an MOI of 10 were used as controls. (A-D) After 5 days of transduction, the cells were subjected to the labeling process with specific antibodies for PD-1 (A) and TIM-3 (B), both conjugated to PE, and the percentage of positive cells was determined by cytometry flow. (C) CD25 expression was verified on M-CAR T cells using flow cytometry. As a positive control for the anti-PD-1, anti-TIM-3, and anti-CD25 labeling step, unmodified Jurkat cells were incubated for 24 h with PMA (50 ng/ml) and PHA–L (5 ug /ml) prior to the marking process for analysis by flow cytometry. The expression of CD25, PD-1, and TIM-3 is identified on the x-axis and the expression of M-CAR is observed on the y-axis, and the histograms (D) represent the fluorescence intensity for labeling with the antibodies mentioned above.

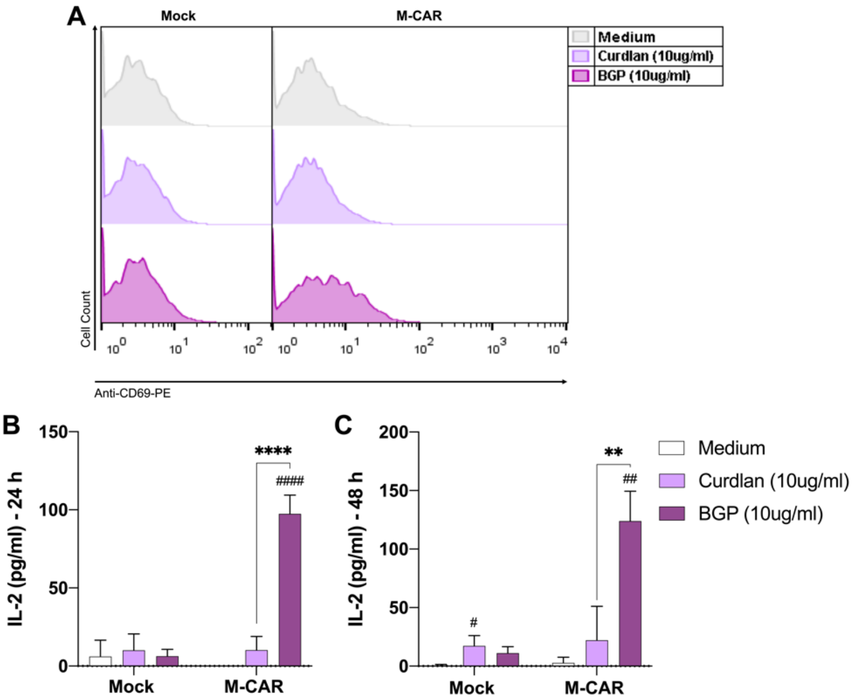


**Supplementary figure 2. Activation of Jurkat M-CAR cells against β-1,3-glucan.** (A) Cell population modified with M-CAR (MOI of 5) and Lenti-mock (MOI of 10) were incubated (2x$10^5$ cells/mL) in 96-well plates in the presence of a β-1,3-glucan polymer, extracted from *Alcaligenes faecalis*, called curdlan (10 ug/ml), or incubated with β-glucan peptide (BGP; 10 ug/ml), extracted from *Trametes versicolor*. M-CAR-mediated cell activation was assessed by CD69 expression after 24 h of culture using flow cytometry. After 24 (B) and 48 h (C) of incubation with 10 ug/ml of curdlan or BGP, cell culture supernatant was obtained for quantification of IL-2 levels by ELISA. The results are expressed as mean ± SD, the significances presented above the bars with "#" are related to the comparison with the mean. #,* p <0.05; ##,** p <0.01; ###,*** p <0.001; ####,**** p<0.0001.

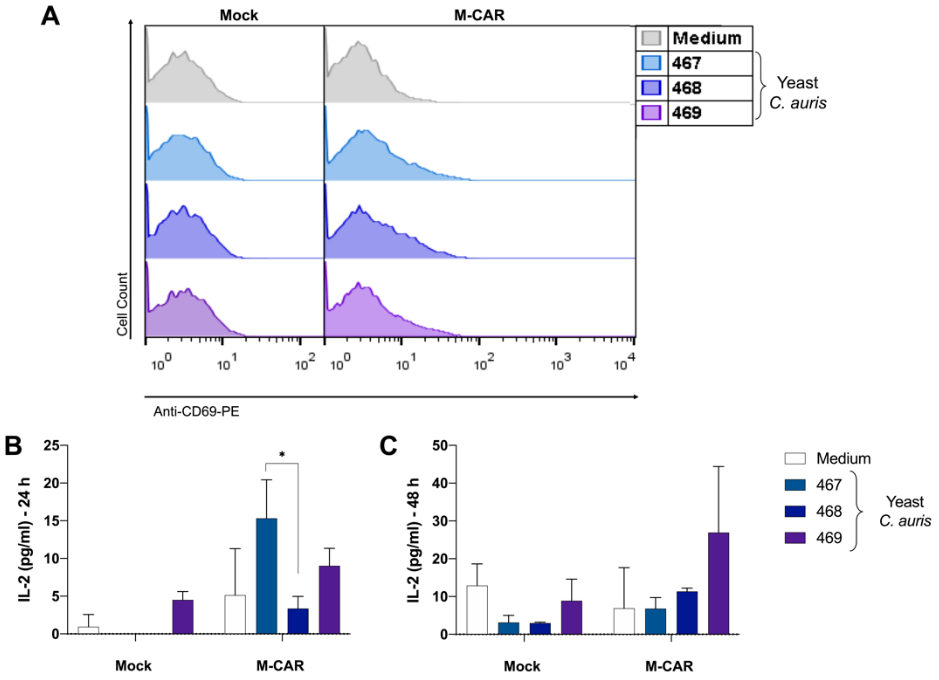


**Supplementary figure 3. CD69 expression and IL-2 production levels in M-CAR cells co-cultured with *C. auris*.** (A-C) Cell population modified with M-CAR (MOI of 5) and Lenti-mock (MOI of 10) were incubated (2x$10^5$ cells/mL) in 96-well plates in the presence of different clinical isolates of *C. auris*, CD69 expression was determined by flow cytometry after 24 h (D), and IL-2 levels in the supernatant of the co-cultivation were quantified after 24 (E) and 48 h (F) by ELISA. The results are expressed as mean ± SD, the significances presented above the bars with "#" are related to the comparison with the mean. #,* p <0.05; ##,** p <0.01; ###,*** p <0.001; ####,**** p<0.0001.

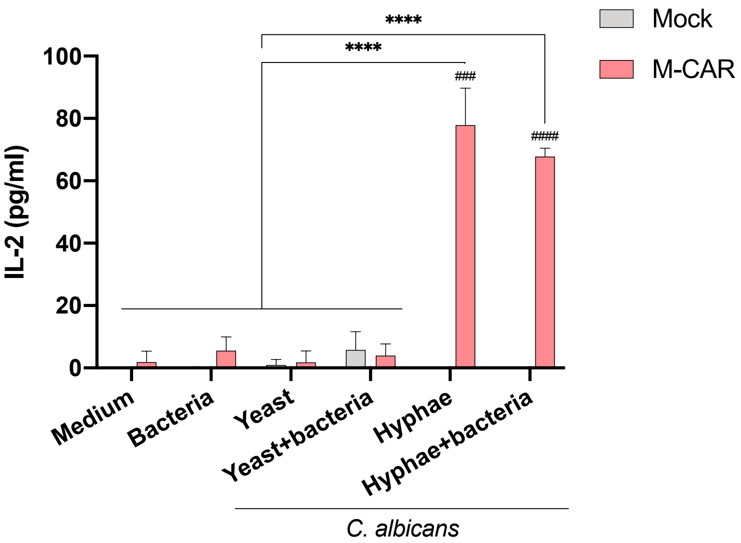


**Supplementary figure 4. Activation of Jurkat M-CAR cells against human microbiota bacteria.** Cell population modified with M-CAR (MOI of 5) and Lenti-mock (MOI of 10) were incubated ($2x10^5$ cells/mL) in 96-well plates in the presence of yeast and hyphae forms of *C.albicans*, and a specific mixture of different bacterial containing *Bifidobacterium breve, B. lactis*, *Lactobacillus paracasei, L. helveticus, L. plantarum, L. acidophilus* and *Streptococcus thermophilus* (ratio of 1:1 cells to bacteria). After 24 h of incubation cell culture supernatant was obtained for quantification of IL-2 levels by ELISA. The results are expressed as mean ± SD, and # represents the significances compared to Lenti-mock. #,*,+ p <0.05; ##,**,++ p <0.01; ###,***,+++ p <0.001; ####,****,++++ p<0.0001.
